# Colibactin DNA damage signature indicates causative role in colorectal cancer

**DOI:** 10.1101/819854

**Authors:** Paulina J. Dziubańska-Kusibab, Hilmar Berger, Federica Battistini, Britta A. M. Bouwman, Amina Iftekhar, Riku Katainen, Nicola Crosetto, Modesto Orozco, Lauri A. Aaltonen, Thomas F. Meyer

## Abstract

Colibactin, a potent genotoxin of *Escherichia coli*, causes DNA double strand breaks (DSBs) in human cells. We investigated if colibactin creates a particular DNA damage signature in infected cells by identifying DSBs in colon cells after infection with *pks*+ *E.coli*. Interestingly, genomic contexts of DSBs were enriched for AT-rich penta-/hexameric sequence motifs, exhibiting a particularly narrow minor groove width and extremely negative electrostatic potential. This corresponded with the binding characteristics of colibactin to double-stranded DNA, as elucidated by docking and molecular dynamics simulations. A survey of somatic mutations at the colibactin target sites of several thousand cancer genomes revealed significant enrichment of the identified motifs in colorectal cancers. Our work provides direct evidence for a role of colibactin in the etiology of human cancer.

**One sentence summary:** We identify a mutational signature of colibactin, which is significantly enriched in human colorectal cancers.

---

The mucosal epithelium is a preferred target of damage by chronic bacterial infections and associated toxins. Not surprisingly, most cancers originate from this tissue. Several infectious agents have been implicated in human cancers, with *Helicobacter pylori* representing the prototype of a cancer-inducing bacterium. Yet, unlike for infections with tumor viruses, which deposit telltale transforming genes in infected cells, for bacterial pathogens compelling evidence of a carcinogenic function is missing due to the lack of specific signatures of past infections in the emerging cancer genomes. Nonetheless, a broader role of bacterial pathogens in human carcinogenesis is highly suggestive.

In humans, several bacterial species have been attributed to a potential role in colorectal cancer (CRC), including *Fusobacterium nucleatum* ^1^and colibactin-producing strains of *E. coli* ^2,3^. Mechanistic analyses indicated distinct cancer-promoting mechanisms elicited by these bacteria, including the activation of inflammatory and growth-promoting signaling pathways as well as the induction of DNA damage ^4^. In particular, colibactin toxin, a secondary metabolite produced by strains of the B2 phylogenetic group of *E. coli*, has long been known to possess DNA damaging ability. In 2006, Nougayrède and collaborators described the 54 kilobase *pks* genomic island that encodes this polyketide-peptide hybrid and showed that *pks*-harboring *E. coli* induce double-strand breaks (DSBs) in host cells and activate the G2-M DNA damage checkpoint pathway ^5^. The recent discovery of a cyclopropane ring, characteristic of DNA alkylating agents, led to the isolation of colibactin-dependent N-3 adenine adducts from host DNA ^6^. This observation was followed by the resolution of colibactin’s mature structure as a highly symmetrical molecule, containing identical cyclopropane warheads at each end, which can give rise to DNA cross-links^7^. Yet, it is unclear if colibactin’s mode of action generates a specific signature that is retrievable in cancers from tissues potentially exposed to respective *E.coli* infections.

To determine a potential preference of colibactin action for specific sites in host cell DNA, we began by globally defining the occurrence of DSBs upon infection of colon derived cells with *pks+ E. coli.* To this end, we applied ‘Breaks Labeling In Situ and Sequencing’ (BLISS), which allows the detection of the exact sites of DSBs in fixed host cells ^8^. The resulting next-generation sequencing (NGS) data and computational analyses revealed a highly specific DNA damage signature, involving AT-rich sequence patterns associated with extreme shape characteristics, which was confirmed by in silico modelling of the colibactin interaction with DNA. By using this information for a stringent search of a mutational signature of colibactin in human cancer genome data, we establish a role of colibactin in the cause of human colorectal cancer and possibly additional cancer types.

## An unbiased sequencing approach to detect colibactin-induced DSB patterns

To confirm colibactin-induced damage, we infected the human colorectal adenocarcinoma cell line Caco-2 with *pks+ E. coli* at MOI 20 for 3 hours. Fluorescence immunohistochemistry showed that cells infected with the wild-type bacteria (*pks+*) were positive for the DNA damage marker γH2AX, while cells infected with the *clb*R deletion mutant (*pks-*), in which colibactin synthesis is restricted ^9^, were not (Fig. 1A). To specifically gain insight into colibactin-induced DSBs and the DNA sequences at which they occur, BLISS was applied to Caco-2 cells infected with WT M1/5 (*pks+*) and mutant Δ*clb*R M1/5 (*pks-*) *E. coli*. Untreated cells and cells treated with the DSB-inducing chemical agent etoposide served as controls (Fig. 1B). BLISS enables unbiased identification of host cell DSBs on a genome-wide scale at nucleotide resolution, based on the amplification of tagged DSBs by *in vitro* transcription. After infection, cells were fixed and the preserved DSBs were blunted *in situ* to allow ligation of specific double-stranded adapters containing a barcode, a unique molecular identifier (UMI), an RA5 Illumina sequencing adapter and a T7 promoter sequence (Fig. 1B). After in vitro transcription, NGS libraries were generated from the produced RNA and sequenced in single-end mode. The included UMIs are used for PCR duplicate removal, while the sample barcodes allow for pooling of different samples prior to the transcription reaction. The raw reads served to determine the genomic positions of DSBs as well as the counts of unique cleavage events using an established analysis pipeline (see Methods). To confirm that our method captured known DSB patterns, we examined the breakpoint density around transcription start sites (TSS), which are reportedly susceptible to breaks induced by etoposide ^8,10,11^. An increase in breakpoint counts around TSSs in our etoposide control was indeed observed (Fig. 1C), indicating the reliability of BLISS as an approach to define colibactin-induced DSB patterns. Next, we performed Locus Overlap Analysis (LOLA) to determine whether the identified DSBs were enriched in particular genomic regions ^12^. Interestingly, unlike the DSBs induced by treatment with etoposide or the DSBs observed in the negative controls, those induced in the *pks*+ *E. coli* condition did not show strong correlation with any known particular genomic regions (Fig. 1D).

**Fig. 1.**
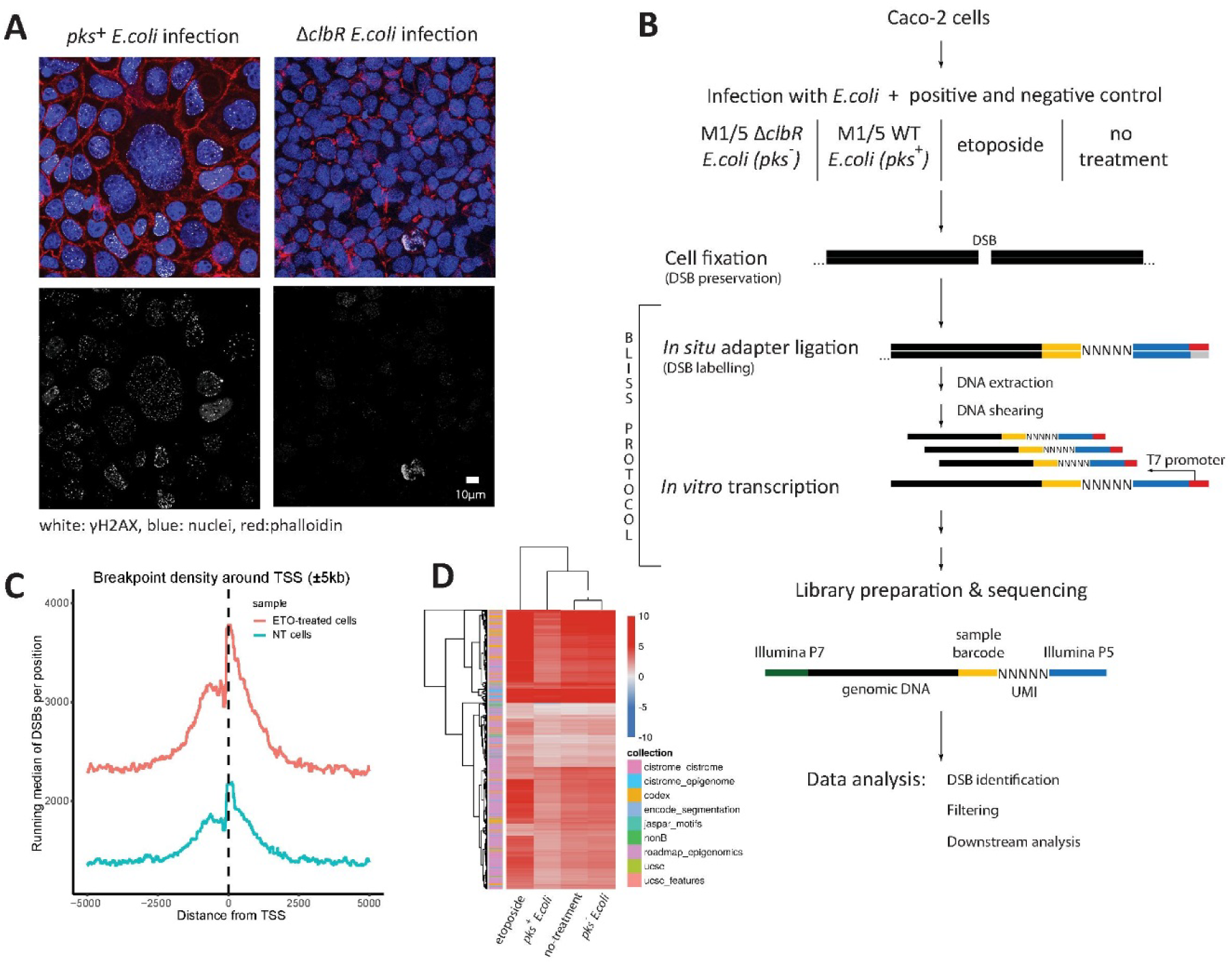
Identification of host DSB upon pks+ E.coli infection. (A) Colibactin-producing E.coli infection causes γH2AX expression in Caco-2 cells. (B) Experimental design for identification of positions of colibactin-induced DSBs with simplified BLISS protocol. (C) BLISS signal of etoposide-induced DSB shows increased counts compared to control condition. (D) Heatmap indicating the log2 odds ratio of break enrichment in genomic region sets (FDR < 5%) compared to the rest of the genome for *pks*+ and *pks*-*E.coli* infected cells, etoposide treated and for non-treated Caco-2 cells.

## Colibactin damages DNA preferentially in specific AT-rich motifs

Next, we asked whether we could identify any particular sequence pattern around the identified DSBs. We thus analyzed nucleotide sequence content of different length stretches around all identified DSBs and compared them between the different treatments. We found that DSBs in cells exposed to *pks*+ *E. coli* are enriched in AT-rich regions. This enrichment was particularly high for the pentanucleotides AAATT and AAAAT together with their complementary mates (Fig. 2A, left panel). This sequence preference of colibactin was evident when compared with either *pks-E. coli* infected or non-treated cells used as the control samples (Fig. S1A). It was detected independently in all four biological replicates, with almost identical relative enrichments (Fig. 2B, Tab. S1). Importantly, no meaningful sequence enrichments were detected when sequence content in close proximity to the DSBs observed in cells exposed to *pks-E. coli* was compared to that in non-treated cells (Fig. 2A, right panel). Hence, the preference for AT-rich sequences is directly linked to the action of colibactin, rather than *E. coli* infection *per se*. To identify the full motif, we analyzed the independent impact of 3’ and 5’ flanking sequences in both identified pentanucleotides for strength of enrichment. Motifs with up to one additional 3’ adenine and/or 5’ thymidine bases were enriched among breakpoints while no impact was observed for more distal nucleotides (Fig. 2C). We also used discriminative motif discovery (DREME)^13^ between breakpoint contexts from *pks+* and *pks-E.coli* infected cells to further narrow down the motif. The top-scoring motif was identified as AAWWTT (Fig. 2D), which contains the enriched pentanucleotide patterns and is compatible with the 3’/5’ extensions represented in Fig. 2C. This symmetric motif indicates a requirement for distant adenines on opposing strands of the double helix while the preference for central A/T nucleotides might derive from dependency on additional conformational conditions.

**Fig. 2.**
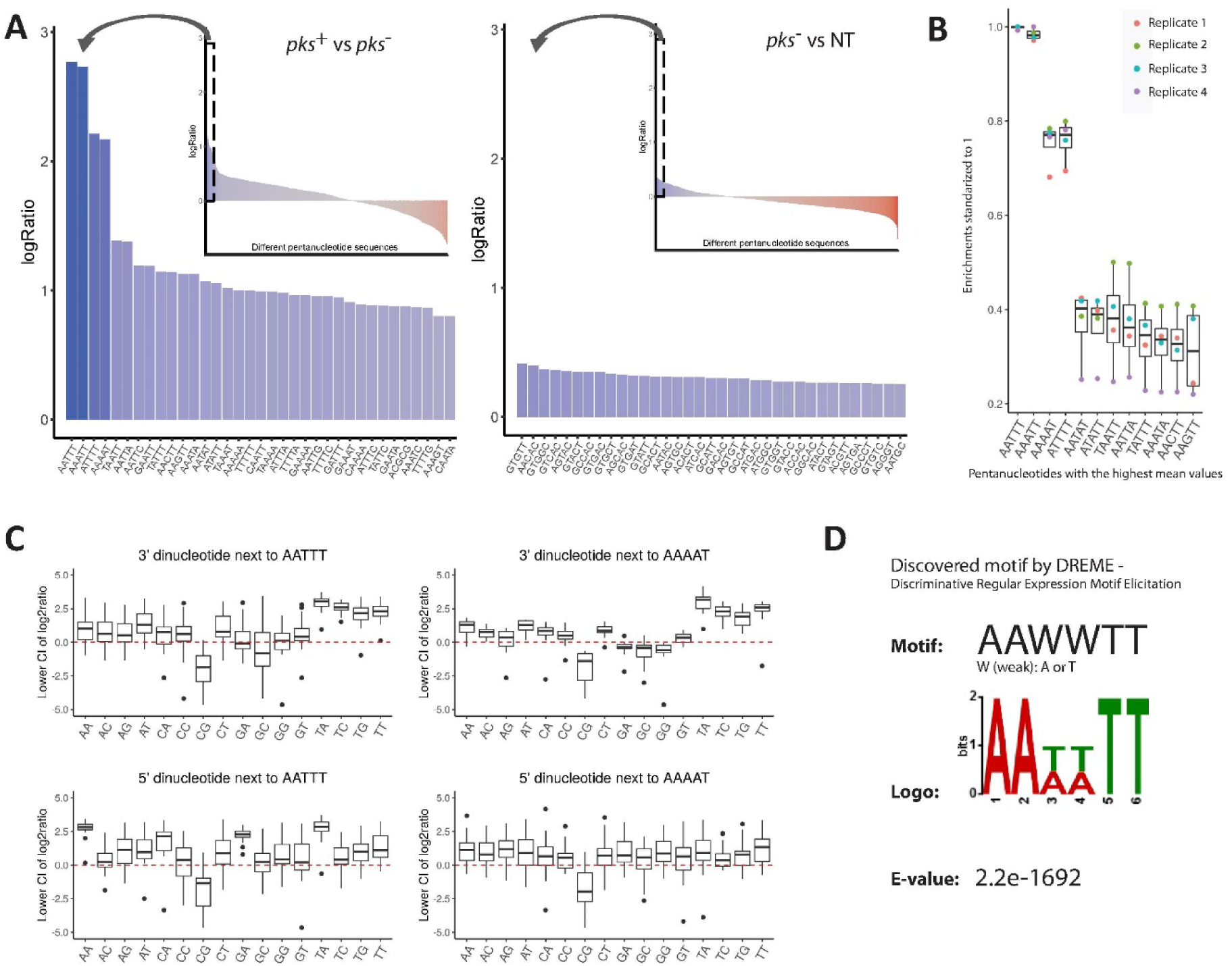
Colibactin damages DNA preferentially in specific AT-rich motifs. (A) Enrichment of pentanucleotide sequences in close proximity to DSB positions (±3 nt) upon different treatments. Plots present pentanucleotide enrichment (log2 ratio of proportions of DSB at each motif between both conditions) of host breaks caused by pks+ E.coli infection in comparison to pks-E.coli infection and caused by pks-E.coli infection in comparison to breaks occurring in the non-treated (NT) cells, respectively. (B) Consistency of an outstanding enrichment of AATTT and ATTTT and their reverse and complement sequences in colibactin induced DSBs in 4 independent biological replicates. Enrichment log2 ratios were standardized so that the highest log2 ratio of each experiment was taken to be 1 and the remaining values scaled accordingly. Enrichments are shown for 11 pentanucleotides with the highest standardized mean values. Each color refers to a different biological replicate. (C) Preferred content of 5’ and 3’ dinucleotides next to colibactin’s pentanucleotids motifs. For each of the motifs (AATTT and AAAAT) we first determined the log2 ratios for all 9nt sequences with the motif in the central 5nt. The 95% confidence interval was computed for each log2 ratio and the distribution of the lower bound of the interval plotted for each possible 2nt sequence at the 5’ or 3’ end of the central pentanucleotide. (D) Top motif enriched in DSBs from pks+ E.coli infected cells compared to DSBs from pks-infected E.coli identified by Discriminative Regular Expression Motif Elicitation (DREME).

## Preferred sites of colibactin action exhibit distinct DNA shape characteristics

Small molecule DNA ligands bind preferentially through intercalation and/or contacts with the double helix major or minor groove, where binding specificity is usually defined by nucleotide sequence-dependent DNA shape characteristics (reviewed by Tse et al.^14^). To investigate whether colibactin has specific DNA shape preferences, we carried out predictions of the shape features in the proximity of each detected DSB. Remarkably, in close proximity (±8 bp) of the detected breakpoints, minor groove width (MGW) exhibited reproducible deviations from the line of averaged values at positions located further away from the DSBs. This was true not only for samples exposed to *pks+ E. coli*, but also for all other samples (Fig. 3A). In addition, all other computationally predicted DNA shape features (helical twist, propeller twist, roll and electrostatic potential) also showed deviations within 8 bp of the DSBs in all samples (Fig. S2). To ensure that the specific landscape of DNA shape at the DSB position is not an artefact of our data analysis approach, the same analysis was performed on 10,000 sequences randomly chosen from the genome (Fig. 3A, inset). Prominent fluctuations of structural properties along the sequences were only observed in close proximity to the identified breaks. Regarding the specific properties of colibactin, we noticed that for all DNA shape patterns the average value at the exact breakpoints in *pks+ E. coli*-infected cells was markedly different from that of all other samples (Fig. 3A enlargement and S2).

**Fig. 3.**
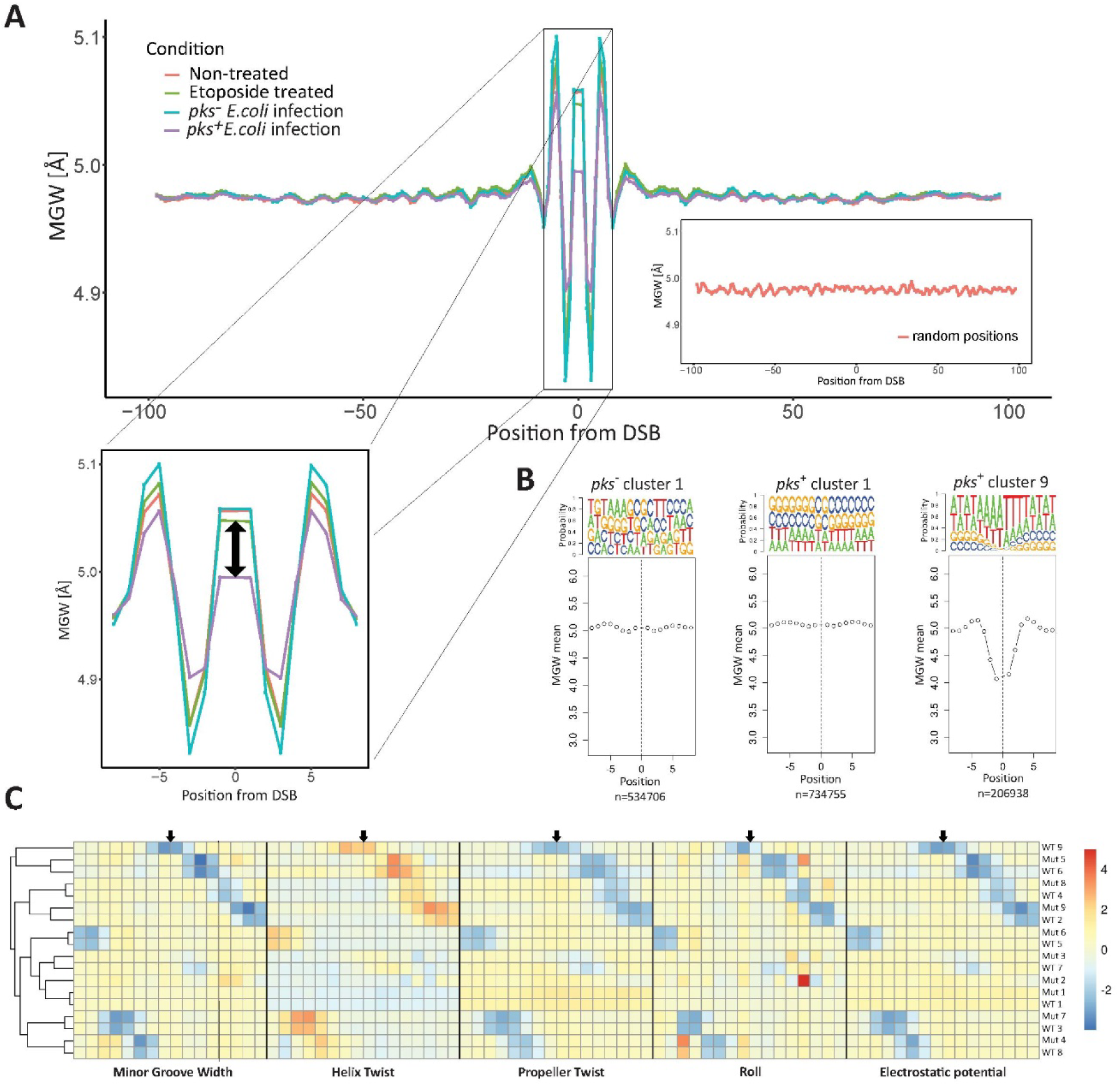
DSB caused by colibactin are associated with specific DNA shape pattern. (A) Averaged minor groove width (MGW) predictions across all +/-100 bp contexts of DSBs identified by BLISS upon different treatments. The difference in averaged profiles of MGW for DSBs between the pks+ E.coli infection condition and all other treatments is enlarged and highlighted by a black arrow. As a control, MGW predictions of flanking sequences of 10,000 randomly chosen genomic positions are presented in the bottom right corner. (B) MGW profiles of selected clusters obtained from k-means clustering of pks+ and pks-conditions. Landscape of cluster 1 for both conditions reflects the general pattern of MGW in close proximity to DSBs (note different y-axis scale). pks+ cluster 9 corresponds to AT-rich sequences across identified DSBs. Profiles of all parameters for every cluster can be found in Supplementary Fig 3A and B. (C) Heatmap comparing averaged profiles of all identified clusters based on all predicted DNA shape parameters across pks+ and pks-infection conditions. Colors indicate individually Z-scored DNA shape characteristics. Each square in the heatmap refers to specific position from the break. Black arrows are marking exact DNA DSB position. Note that pks+ cluster 9 is unique for this treatment and shows extreme values centered at the DSB position.

Averaged profiles describe the superposition of potentially many underlying shape motifs, many of which might be attributable to DSBs generated by processes other than colibactin and are shared across conditions. To further explore the differences between the DNA shape parameters in the individual breakpoint positions of *pks+* and *pks-E. coli* infections in an unbiased manner, we applied k-means clustering as unsupervised machine learning algorithm. Assigning every set of predicted values of the DNA shape characteristics for each DSB to the closest centroid of 1 out of 9 clusters independently for both *pks*_+_ and *pks*_−_ *E. coli* induced DSBs, resulted in specific and unique shape patterns for each cluster (Fig. S3A,B). Interestingly, a quarter of all breakpoints from both infection models were assigned to the respective cluster 1 (Fig. 3B), whose profile amplitudes and pattern correspond to the global profile of MGW. To gain a better overview of the sequence content of each cluster, the probability for the presence of each nucleotide was computed for each position (top row for each cluster). As expected, MGW dips were associated with high AT-rich content in all clusters. Most of those dips correlate to short AT stretches most likely caused by periodic 10 bp spaced WW dinucleotide motifs in the genome sequence associated with nucleosome positioning^15^, which occur at different positions in the breakpoint context and are therefore distributed to separate clusters.

To determine whether any cluster was unique to a particular treatment, we compared clusters for both infection conditions. Indeed, cluster 9 of the *pks+ E. coli* dataset was not paired with any *pks-*cluster for any of the parameters examined and showed the strongest deviations of shape parameters centered at the estimated DSB position. This was true regardless of whether clusters were compared separately for each predicted DNA shape parameter (Fig. S3C) or for all parameters together (Fig. 3C). This confirms that the sequences in proximity to the DSBs assigned to cluster 9 of *pks+ E. coli* represent a group of breaks unique to this condition. Differences between MGW means for cluster 1 and 9 show how strongly AT-rich sequences influence local DNA shape (Fig. 3B).

## Colibactin’s binding motif corresponds to extreme DNA shape parameter values

In order to unveil the features of the DNA molecules preferred by colibactin, we analyzed the structural properties of the DNA stretches close to the identified DSBs. We correlated the predicted DNA shape parameters for the central 1-2 bps of all possible pentanucleotides (1024) with the log2-ratios of pentanucleotide sequence enrichment in DSB positions caused by colibactin. Remarkably, colibactin’s preferred pentanucleotide sequences, d(AAATT)·d(AATTT) and d(AAAAT)·d(ATTTT), were associated with the narrowest minor groove widths, with values below 3 and 3.7 Å respectively, as well as with some of the most negative values for propeller twist of the central base pair and extremely negative electrostatic potential (Fig. 4A). Closer inspection of the inter-base pair parameter roll revealed that AAATT, the most frequent pentamer that surrounds the break point, also shows very peculiar conformational characteristics (Figure 4A). The DNA stretch composed of A-tract followed by T-tract tract shows that the progressive narrowing of the DNA minor groove going from the 5’ to the 3’ end is correlated with low roll values. Values for the DNA stiffness descriptor (40) (k_tot, see Methods section), revealed that these tracts possess high intrinsic rigidity (Fig. 4A, Table S1), making them difficult to distort.

**Fig. 4.**
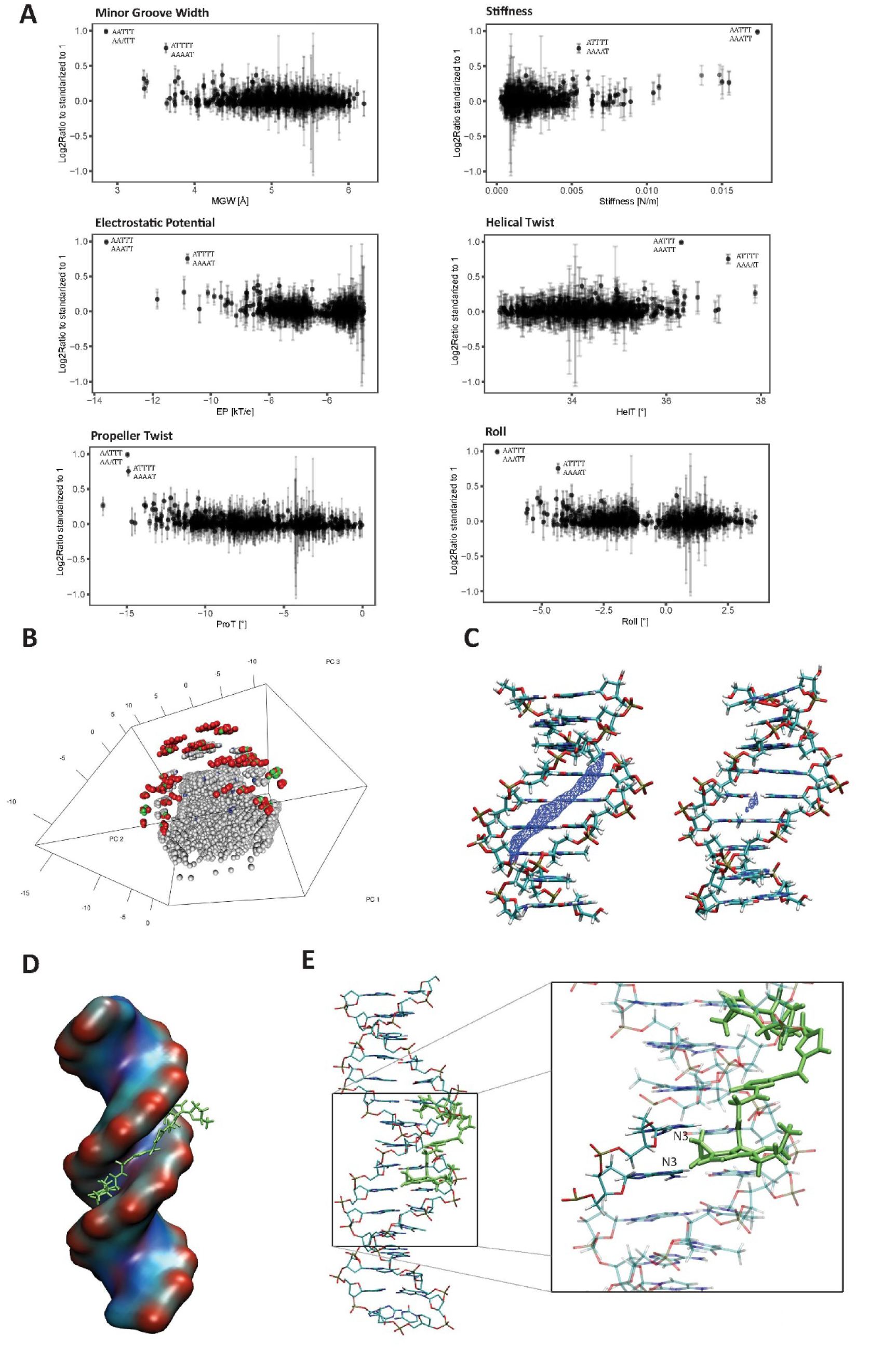
Colibactin’s binding motif corresponds to extreme DNA shape parameters values and extreme value of electrostatic potential. (A) Correlation between pentanucleotide sequence enrichments (standardized to 1; for 4 biological replicates) for colibactin’s activity and values of predicted DNA shape parameters. For MGW and EP values are calculated for each pentamer; for ProT intra-base pair parameter for the central base pair of each pentamer are calculated; for Roll and HelT the average of the two inter-base pair parameters, considering the two central base pair steps in each pentamer is calculated; for Stiffness average values, considering the two central base pair steps in each pentamer are calculated. (B) 3D visualization of the first 3 principal components from predicted DNA shape values for the central 5nt of all possible 9nt motifs. Those 9nt motifs containing AAWWTT and/or showing strong enrichment are highlighted. Labels: red – AAWWTT motif with lower 95% confidence interval (CI) limit of log2ratio > 1.5; green – AAWWTT with lower CI limit < 1.5; blue – non-AAWWTT sequences with lower CI limit > 1.5, grey – other sequences (proportionally downsampled to approximately 35,000 sequences). (C) Molecular Interaction Potential (MIP) using Na^+^ as probe for 2 cases, on the left the most preferred 9nt DNA sequence for colibactin binding (CAAATTTTG) and on the right the least favorite (AACTTTGCA). The isosurfaces (in blue) for the two DNA sequences show different electrostatic potential (isovalue =-7 kcal mol^-1^), correlating with the different minor groove conformations. (D-E) Images of the theoretical docking of predicted colibactin structure into its preferred sequence motif (central sequence AAATTT), showing the insertion of the colibactin into the minor groove, with the double stranded DNA as surface (D) and showing the atomic details (E). Enlargement shows a zoomed-in image of the closeness of the cyclopropane to one of the N3 atom of the adenine, highlighting the possibility to alkylate the consequential base pair depending on the carbon involved in the alkylation.

To obtain a more complete picture of the combined effect of the DNA shape characteristics, we extended this analysis to the central 5 bp of all possible 9 bp sequences and explored the multivariate space defined by all DNA shape parameters and at all positions by principal component analysis. Again, enriched motifs in pks+ E. coli infected cells compared to pks-E. coli stood out as an extreme group among all analyzed sequences (Fig. 4B). The data suggest that colibactin’s binding preference for DNA stretches with the central pentanucleotides AAATT/AAAAT is driven not only by nucleotide content but also by particularly extreme values of sequence-associated DNA shape attributes like MGW and electrostatic potential. To probe this, we also calculated the molecular interaction potential (MIP) using Na^+^ as probe for the most and the least preferred DNA central pentamers for colibactin binding (AATTT and CTTTG respectively). The isosurfaces for the two DNA sequences (Figure 4C, blue) confirmed strongly different electrostatic potential correlated with different minor groove conformations, which is likely to be related to the difference in colibactin binding affinity. All these observations suggest that the unusually narrow minor groove together with an inherent rigidity and a marked electrostatic potential facilitate recognition and binding of colibactin, probably maximizing its interactions with the DNA.

In order to explore the binding between the DNA and colibactin we built a molecular model of colibactin (see Methods for details) using quantum mechanics (QM) calculations as first structural guess. The optimized structures were then hydrated and subjected to molecular dynamics (MD) simulations (details on parametrization are discussed in Methods) using state-of-the-art simulation conditions (see Methods). Colibactin appears as a rather flexible molecule, with an average end-to-end distance around 13 Å (Fig. S4). This suggests it can bind 4-5 base pairs if located along the minor groove, which is supported by its structure, its preference for AT-rich sequences, and its ability to attack N3. HADDOCK software ^16^ was used as docking engine, to obtain a putative binding mode. The default scoring function was supplemented by restraints forcing the orientation of the reactive cyclopropane moiety towards the N3 of the adenine. The best docking poses were manually curated and subjected to MD simulations (see Methods). The final putative model shows a very stable binding of colibactin to the minor groove (Figure 4D), with excellent van der Waals contacts with all the walls of the groove and the cyclopropane rings pointing towards the adenines on opposite strands (Fig. 4E). From the equilibrium trajectory we determined that the number of base pairs involved in the binding could fluctuate between 4 and 5, depending on the orientation of the cyclopropane, and the carbon alkylating the N3 of the adenines (Figure 4E, enlargement). In all cases colibactin fits perfectly into the narrow minor groove of the targeted sequences and adopts a spatial arrangement that would facility alkylation at N3.

## Somatic mutations at colibactin target sequences indicate role in cancerogenesis

Having identified a specific nucleotide sequence associated with colibactin-induced DSBs, we wondered if we could identify a specific mutational signature associated with this sequence in cancers that have been experimentally connected to *pks+ E. coli* infection ^2,17^. Using whole-exome sequencing (WXS) data from colorectal cancer samples ^18^ (n=619) and across several cancer entities in the TCGA project (https://www.cancer.gov/tcga, see Methods, n=553 colorectal cancers among 10,224 tumor cases in 24 cancer types), we tested whether somatic mutations are specifically enriched at the identified pentanucleotide sequences. We determined the hexanucleotide-specific mutation rate for all possible hexanucleotides adjusted for their frequency in exonic regions. Given colibactin’s demonstrated preference for alkylation of adenines, we assessed the mutation rate for single nucleotide variants (SNV) at reference bases A or T. We hypothesized that preferential binding of colibactin to AAWWTT motifs (i.e. AAATTT or AATTTT/AAAATT) should increase the mutation rate at these motifs compared to all other hexanucleotides with the same length and nucleotide content (i.e. all remaining WWWWWW motifs). Since we observed that mutation rates at AAWWTT motifs were particularly high in hypermutator samples harbouring polymerase epsilon (POLE) mutations, we assessed mutation rates in cohorts defined by total SNV numbers per samples and POLE-mutated samples separately. We found that mutation rates in AAWWTT motifs were enriched compared to all other WWWWWW motifs in colorectal cancers in both data sets analyzed (Fig. 5A). In the TCGA pan-cancer data set we also found enrichment at AAWWTT motifs in stomach cancer, uterine corpus endometroid cancer and breast cancer. No enrichment was found, e.g. in head and neck squamous cancer, lung adenocarcinoma and lung squamous carcinoma, while enrichment only for POLE mutated cases was found in bladder cancer and cervical squamous cancer (Fig. 5B).

**Fig. 5.**
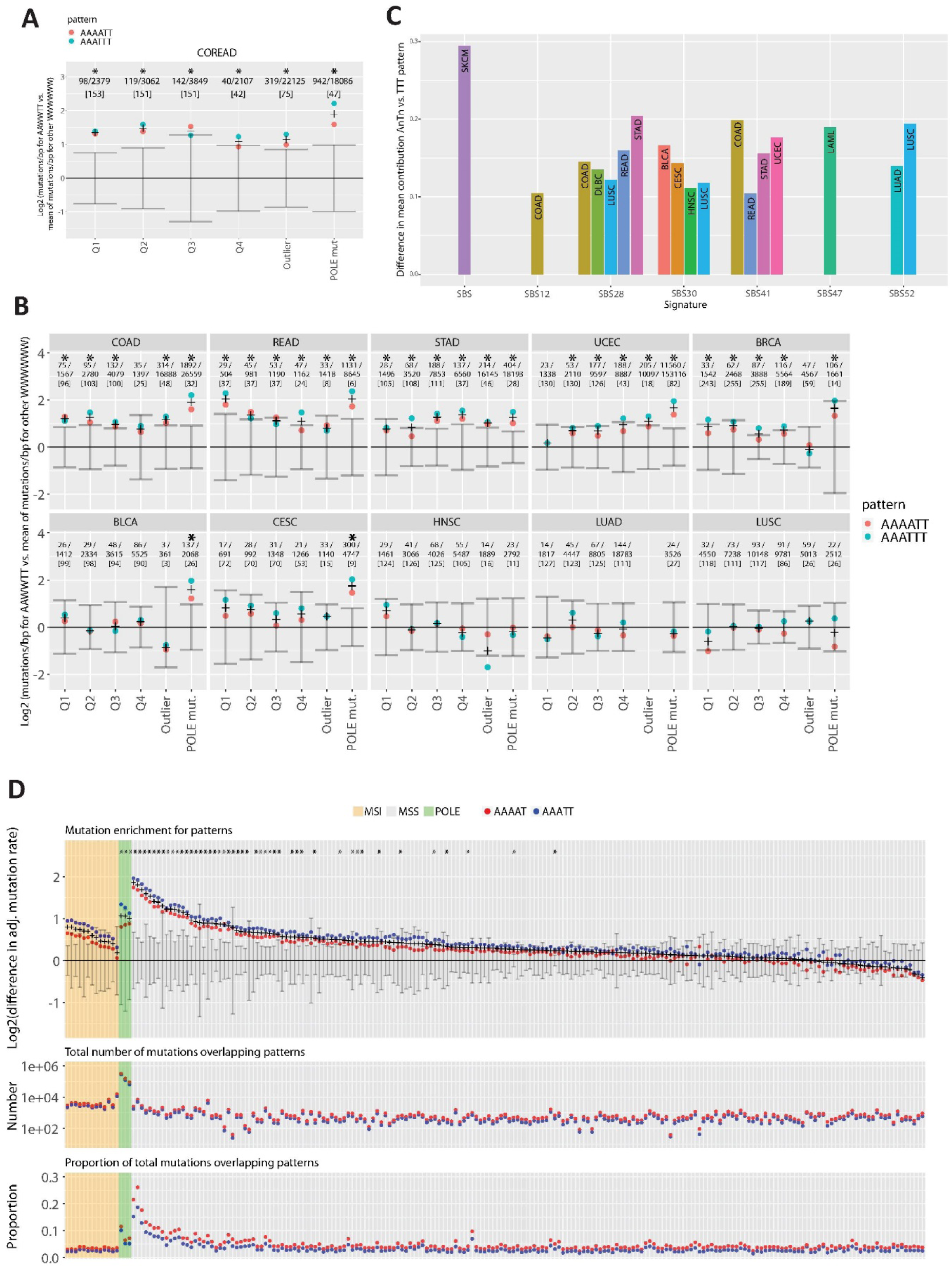
Several cancers show enrichment of mutations at colbactin associated motifs. A) Enrichment of single base change (SBS) mutations at colibactin-associated hexanucleotide motifs AAATTT/AAAATT in exome sequences from colorectal cancer cases ^18^ B) Enrichment of SBS mutations at colibactin associated hexanucleotide motifs AAATTT/AAAATT in exome sequences from TCGA. Top row: cancer entities showing enrichment across all subcohorts. Bottom row: cancer entities showing enrichment only for POLE mutated cases or no enrichment at all. COAD-colon adenocarcinoma, READ-rectal adenocarcinoma, STAD-stomach adenocarcinoma, UCEC-uterine corpus endometroid cancer, BRCA-breast cancer, BLCA-bladder cancer, CESC-cervix squamous cell carcinoma, HNSC-head and neck squamous cell cancer, LUAD-lung adenomcarcinoma, LUSC-lung squamous cell carcinoma C) Signature detection rates for SBS mutations with contexts overlapping AATTT/ATTTT. Only signatures with significant and positive differences in signature detection rates for contexts overlapping AATTT/ATTTT compared to TTT are shown. D) Analysis of SBS mutations at colibactin associated pentanucleotide motifs AAATT/AAAAT in whole genome somatic mutation data from ^19^. Top: Difference in log2(mutations/bp covered by motif) between colibactin associated and all other WWWWW motifs. Middle: Total mutation count at colibactin associated motifs: Bottom: Proportion of total mutations ovlerapping colibactin associated motifs. MSS,MSI, and POLE, mutated cases. (A), (B), (D) Stars denote significant difference (Mann-Whitney-U test p < 0.05 and FDR < 20%) between colibactin associated motifs and all other motifs with the same A:T content and length (A, B: hexanucleotide (HN) motif: AAATTT/AAAATT vs WWWWWW motifs, C: pentanucleotide (PN) motifs: AAATT/AAAAT vs. WWWWW motif). (A,B): First line is [number of mutations overlapping AAWWTT motif] / [all mutations in cohort]. Third lines is number of samples in cohort. Error bars describe the ± 2MAD intervals for mutation rate (mutations/bp covered by motif) of WWWWW(W) motifs excluding colbactin associated pattern after subtracting their mean. Dots represent the mutation rates for the two colibactin associated PN or HN motifs after subtracting the mean of the WWWWW(W) motifs. Crosses are the mean of the colibactin associated motifs.

We validated the findings from WXS data in a cohort of colorectal cancer assessed by whole genome sequencing (WGS) ^19^. We analyzed enrichment of mutations at colibactin associated motifs for 208 tumors including 193 microsatellite stable (MSS), 3 POLE mutated and 12 microsatellite instable (MSI) cases in a similar way as for WXS data but considering each sample separately instead of pooling in subcohorts. This allowed to identify enrichment and mutational loads for individual samples. We found significant (Mann-Whitney-U test, p<0.05, FDR <20%) enrichment of mutations at colibactin associated pentanucleotide motifs compared to other motifs with same length and A/T content in 3/3 POLE mutated samples and 49/193 (25.3%) MSS cases but not in MSI cases. We found similar enrichment as for penta-(AAATT/AAAAT) for hexanucleotide (AAWWTT) motifs associated with colibactin in MSS samples (data not shown). The median number of mutations in MSS samples at colibactin associated motifs was 963 (range: 63-11876) corresponding to a median proportion of 6.7% (range: 3.9-44.7%).

We next asked if an association exists between the preferred colibactin motif and any of the previously described mutational signatures^20,21^. Again, we used somatic mutation data from the TCGA data set as above and classified all single nucleotide variants according to the sequence context in direct proximity (+/− 5bp). Variants were assigned to one of three groups: Those with sequence context containing AAATT/AATTT or AAAAT/ATTTT, those with contexts containing a control TTT motif and all remaining mutations. Globally we observed distinct mutation frequencies for several trinucleotide changes (Fig. 5C) in those classes. We identified a contribution of known signatures in those 3 classes for all samples and selected those with significantly higher contributions at AATTT or AAAT motifs compared to TTT and all other motifs (Fig. 5D). Two of the signatures with increased contributions at colibactin-associated motifs where of particular interest: SBS28 (https://cancer.sanger.ac.uk/cosmic/signatures/SBS/SBS28.tt) and SBS41 (https://cancer.sanger.ac.uk/cosmic/signatures/SBS/SBS41.tt) both with unknown etiology and featuring predominantly mutations at T:A and a prominent T[T>G]T trinucleotide change, were found enriched in colorectal cancers, among others. While SBS28 has been previously shown to be associated with POLE mutation-related hypermutated tumors, SBS41 was enriched in stomach adenocarcinoma, colorectal adenocarcinoma and endometrial carcinoma of the uterine corpus, mirroring the results for motif enrichment above.

## Discussion

We pursued an unbiased bimodal approach that revealed a signature of the bacterial genotoxin colibactin in the human cancer genome indicating a causal link between a bacterial infection and the emergence of cancer. This was achieved by first defining the DSB-landscape generated by the action of colibactin through applying the BLISS sequencing technology and subsequent comprehensive analysis of the genome-wide location of DSBs. The resulting DSB pattern, which exhibits exceptional structural features, corresponded to, and could be further refined by, three-dimensional modelling of the colibactin–DNA complex, involving distinct topological interactions with the minor-groove. In a second step, we used the identified motif to assign associated mutations in various cancer genome databases. Most interestingly, we revealed an enrichment of mutations at colibactin-associated motifs in colorectal cancers but also detectable in a few other cancer types, notably uterine endometroid and stomach cancer. We identified putative trinucleotide signatures (SBS41, SBS28) in the context of these mutant sites in the same cancer entities.

The identified AAATT and AAAAT motifs are associated with extreme physical values of the DNA duplex, most prominently characterized by a very narrow minor groove width, which generates highly negative electrostatic potential and renders the DNA segment stiff. This extreme physical property implicates a low propensity of the colibactin target site to bind to proteins ^22^. In fact, poly(dA-dT)-tracts are rarely found inside nucleosomes, but are prevalent in nucleosome-free regions (NFRs) ^22^. Thus, colibactin’s particular targeting preferences for non-protected DNA regions might increase the efficacy of the toxin. Even though definitive evidence of the binding conformation requires further experimental support from 3D structural analysis, the 3D model provided here allows for demonstrating and validating the extraordinary electrostatic properties of the identified motifs and the fit of colibactin to the minor groove. It puts a limit of 4-5 nucleotides on the distance of adenines attacked by the cyclopropane groups of the same molecule. Although most of the DNA shape characteristics are directly driven by the underlying sequence, the fact that other sequences with similar A/T content were not strongly enriched around DSBs indicates a dominance of DNA shape over sequence characteristics for the binding of colibactin.

Similar DNA shape and sequence affinities have been reported for other bacterial DNA toxins, such as duocarmycin, yatakemycin, distamycin, netropsin and CC-1065 – small molecules produced by *Streptomyces* spp., which are all minor groove binders with AT-rich sequence selectivity. Distamycin ^23^ and netropsin ^24^ act as RNA and DNA polymerase inhibitors ^25^, while duocarmycin, yatakemycin and CC-1065 are DNA alkylators. Duocarmycin ^26^ selectively alkylates adenine residues flanked by three 5’-A or T-bases (5’-WWWA-3’) ^27^, yatakemycin ^28^ preferentially alkylates the central adenine of a five-base AT site (5’-WWAWW-3’) ^29^ and CC-1065 ^30,31^ shows selectivity for more extended five-base AT-rich alkylation sites (5’-WWWWA-3’) ^27^. The fact that these toxins possess similar mechanisms of action, even though they derive from different bacterial strains, suggests that they arose via convergent evolution. Genotoxins are widespread amongst bacterial species, where they are thought to serve primarily for inter-microbial competition ^32^. Unsurprisingly, therefore, all of the mentioned alkylating toxins inhibit the growth of many Gram-positive and Gram-negative bacteria as well as some pathogenic fungi, such as *Aspergillus fumigatus* and *Candida albicans* ^28,30^. Similarly, *pks+ E.coli* inhibit the growth of *Staphylococcus aureus*, also in its multi-resistant form ^33^.

How colibactin-induced DNA damage is repaired is still unknown. Different host DNA repair mechanisms can be involved depending possibly also on the cell cycle phase. Effects of repair involve nucleotide excision ^34^ of alkylated adenines which could lead to DSBs, resection of break ends or complete repair, or error-prone repair by translesion DNA polymerases in late phases of the cell cycle, among others. We were able to show enrichment of SNV at colibactin-associated motifs in exome and whole-genome sequencing datasets. For colorectal cancers, whole-genome sequences revealed elevated mutation rates in colibactin associated motifs in at least 25% of all MSS cases and a colibactin attributable mutation load of around 6% in most patients. Further analyses of whole-genome sequenced samples including the analysis of breakpoints of structural variants will be required to assess the full spectrum of damage-related mutations in host cells. Mutational signatures for other alkylating substances, such as cisplatin, have been identified in human DNA sequences after exposure to the mutagen ^21,35^. However, it is to be expected that the signatures depend strongly on the specific type of damage induced by each substance. Here we identified two signatures that are consistent with colibactin action, one with (SBS28) and one without (SBS41) relation to known DNA repair defects. An impact of reduced DNA repair and mutagen-induced damage on the emergence of different mutational signatures has recently been shown in a model of *C. elegans* ^36^. The enrichment of mutations specifically in POLE cases hints at either a similar outcome of distinct mutational processes or even a role of POLE in the repair of colibactin-associated damage.

Colibactin has been found not only in *E. coli* but also in *Klebsiella* isolates ^37^. Considering the widespread and diversity of bacteria carrying this toxin, it is maybe not surprising that the mutational signature identified here is not only restricted to the colon. Rather, other tissues might also be colonized by either *pks*+ E. coli, another species bearing the *pks* gene cluster, or a different species with a closely related genotoxin. Thus, our study will stimulate future research on other pathogen-host cell encounters that could lead to an even greater match of the identified signature with different cancer types. Better understanding of the role of the microbiome in malignant degeneration should provide new and exciting opportunities for cancer prevention.

## Materials and Methods

### Cell line, bacterial strains, E.coli infection and etoposide-treatment

Caco-2 cells (from ATCC^®^ HTB-37™) were cultured at 37 °C under a water-saturated 5% CO_2_ atmosphere, in DMEM medium (Life Technologies, cat. number: 10938-025), supplemented with 20% FCS (Biochrom, cat. number: S0115). Contamination of Mycoplasma spp. in immortal cell line was excluded using Venor^®^GeM OneStep PCR kit (Minerva Biolabs^®^, cat. number: 11-8250). To infect Caco-2 cells, overnight liquid culture of E.coli strain M1/5 (Streptomycin-resistant and colibactin-positive) and E.coli strain M1/5::ΔclbR (streptomycin-resistant and colibactin-negative) was set up. Bacteria were inoculated in 5 ml of Luria broth (LB) medium and incubated overnight at 37 °C in a shaking incubator. The overnight inoculum was diluted 1:33 in infection medium (DMEM + 10% FCS + HEPES (Life Technologies, cat. number: 15630-056)) to obtain OD_600_=1 after 3 h of incubation to give 1.5 × 10^9^ bacteria/ml. Prepared bacteria inoculum was further diluted to reach MOI 20, added to Caco-2 cells seeded previously and incubated for 3 hours at 37 °C. Medium was then aspirated and cells fixed according to the protocol for immunofluorescence or BLISS. For every biological replicate positive (etoposide-treatment) and negative (no treatment) controls were included. Etoposide powder (Sigma Aldrich, cat. number: E1383) was diluted in DMSO in order to reach 50 mM working solution. Aliquots of the drug were stored at −20 °C. Final drug dilutions to the concentration of 50 μM were performed in pre-warmed infection medium prior to each drug exposure. Treatment was conducted for 3 hours at 37 °C and afterwards medium was aspirated and etoposide-treated cells were fixed in the same way as E. coli-infected cells.

### Immunofluorescence staining

Caco-2 cells grown and infected on MatTek glass-bottom dishes were washed three times with PBS (Life Technologies, cat. number: 14190-094) and fixed with 3.7% paraformaldehyde (Sigma Aldrich, cat. number: P6148) for 1 h. The cells were kept overnight in blocking buffer (3% BSA, Biomol, cat. number: 01400.100), 1% saponin (Sigma Aldrich, cat. number: 84510), 2% Triton X-100 (Carl Roth, cat. number: 3051.2) and 0.02% sodium azide (Sigma Aldrich, cat. number: S2002). Blocking was followed by overnight incubation with γH2AX antibody (Phospho-Histone H2A.X (Ser139) Antibody, Cell Signaling, cat. number: 2577, 1:500 dilution) at 4 °C. The next day, the MatTek dishes were washed three times with blocking buffer followed by overnight incubation with secondary antibody (Dianova, cat. number: 711-035-152, 1:250 dilution) diluted in blocking buffer. Phalloidin 546 (Invitrogen, cat. number: A22283, 1:200 dilution) and Hoechst (Sigma, cat. number: H6024, 1:10000 dilution) were added for staining actin filaments and DNA, respectively. The next day, cells were washed three times with blocking buffer and coverslipped using Vectashield^®^ Antifade Mounting Medium (Vector Laboratories, cat. number: H-1000). Images were acquired using a Leica TCS SP-8 confocal microscope and processed using ImageJ.

### sBLISS, an adaptation of the BLISS method

DSBs were identified using the suspension-cell BLISS (sBLISS) method ^38^, which is an adaptation of the previously published BLISS protocol ^8,39^. In contrast to BLISS, where DSBs are labeled in fixed cells immobilized on microscope slide, in sBLISS DSBs are labeled in fixed cell suspensions. In brief, cells were treated/infected in culture dishes and afterwards trypsinized, counted, centrifuged and resuspended in pre-warmed medium to obtain 10^6^ cells per 1 ml. Then, cells were fixed by adding 16% PFA (Electron Microscopy Sciences, cat. number: 15710) to reach a final concentration of 4%. After 10 minutes, 2 M glycine (Molecular Dimensions, cat. number: MD2-100-105) was added to a final concentration of 125 mM in order to block unreacted aldehydes. This was followed by two 5 minutes incubations, first at room temperature and then on ice, followed by two washes in ice-cold PBS. Cross-linked cells were stored in PBS at 4 °C until further processing.

Next, BLISS template was prepared. This includes: (1) Cell lysis in 10mM Tris-HCl, 10 mM NaCl, 1 mM EDTA, and 0.2% Triton X-100 (pH 8) buffer, followed with lysis in buffer containing 10 mM Tris-HCl, 150 mM NaCl, 1 mM EDTA, and 0.3% SDS (pH 8); (2) DSBs blunting with NEB’s Quick Blunting Kit (NEB, cat. number: E1201); (3) *In situ* BLISS adapter ligation using T4 DNA Ligase (ThermoFisher Scientific, cat. number: EL0011). Each BLISS adapter contained a T7 promoter sequence for IVT, the RA5 Illumina RNA adapter sequence, a random 8nt long sequence referred to as Unique Molecular Identifier (UMI) and a 8nt long sample barcode; (4) Phenol:chloroform-based extraction of gDNA; (5) Fragmentation of isolated genomic DNA (400-600bp) using BioRuptor Plus (Diagenode). Obtained BLISS templates were stored at −20 °C.

The final step of the BLISS protocol was *in vitro* transcription (IVT) followed by NGS library preparation. At first, 100ng of purified, sonicated and differentially-barcoded BLISS template of 1) etoposide-treated and non-treated cells, or 2) cells infected with pks+ E.coli or infected with pks-E.coli were pooled into one reaction, respectively. IVT was performed using MEGAscript T7 Transcription Kit (ThermoFisher, cat. number: AMB13345) for 14 hours at 37 °C in the presence of RiboSafe RNAse Inhibitor (Bioline, cat. number BIO-65028). Next, gDNA was removed using DNase I (ThermoFisher, cat. number: AM2222) and the remaining RNA was purified with Agencourt RNAClean XP beads (Beckman Coulter). The Illumina RA3 adapter sequence was ligated to the purified RNA using T4 RNA Ligase 2 (NEB, cat. number: M0242) for 2 hours at 25 °C and reverse transcription was performed with Reverse Transcription Primer (Illumina sequence) using SuperScript IV Reverse Transcriptase (ThermoFisher, cat. number: 18090050) for 50 minutes at 50 °C. This was followed by enzyme heat inactivation for 10 minutes at 80 °C. Finally, libraries were amplified with NEBNext High-Fidelty 2x PCR Master Mix (NEB, cat. number: M0541), the RP1 common primer and a uniquely selected index primer. 12 PCR cycles were conducted, and after that libraries were purified according to the two-sided AMPure XP bead purification protocol (Beckman Coulter). Profiles of the libraries were quantified on a BioAnalyzer High Sensitivity DNA chip. Libraries were sequenced as single-end (1×75) reads on the NextSeq platform.

### Pre-processing of sequencing data

Raw sequencing data were pre-processed as previously described ^7^. In brief, only reads which contained the expected prefix of UMI and sample barcode were kept using SAMtools ^40^. One mismatch in the barcode sequence was allowed. Further, prefixes were trimmed and the remaining sequences were aligned to the GRCh37/hg19 reference genome using BWA-MEM ^41^. Reads with mapping quality scores ≤ 30 and those which were determined as PCR duplicates were removed. Finally, a BED file containing a list of unique DSBs locations was generated. DSBs which fell into ENCODE blacklist regions ^42^, high coverage regions ^34^ and low mappability regions ^34^ were removed. Kept positions of DSBs were further used in downstream analysis.

### Locus Overlap Analysis

To identify significant overlaps of DNA DSB with genomic region sets we used LOLA ^11^. We first defined whole genome as a Universe Set, which was next divided into tiles of equal lengths (1,000 nt). For each created tile we next searched for overlaps with captured by BLISS DSBs using the findOverlap() function. All tiles containing ≥ 10 breaks were used as a Query Set. The runLOLA() function was executed with LOLA Core databases (reduced by Tissue clustered DNase hypersensitive sites) as well as LOLA Extended databases and custom database containing non-B-DNA regions (https://nonb-abcc.ncifcrf.gov/apps/site/references). Fisher’s exact test was used with a FDR ≤ 5%.

### DNA Shape predictions

DNA structures can be described in terms of base-pair and base-step parameters that consist of three translational and rotational movements between the bases or the base pairs, respectively. At the base-pair step level, DNA deformability along these six directions has been described by the associated stiffness matrix ^43^. From the ensemble of MD simulations considering the tetramer environment using the newly refined parmbsc1 force field, we retrieved the 6×6 matrix describing the deformability of the helical parameters for each possible DNA tetramer. Pure stiffness constants corresponding to the six base-pair step parameters (shift, slide, rise, tilt, roll and twist) were extracted from the diagonal of the matrix and the total stiffness (*K_*tot) was obtained as a product of these six constants and used as an estimate of the flexibility of each base pair step in a tetramer. For predictions of minor groove width (MGW), propeller twist (ProT), electrostatic potential (EP), helical twist (HelT) and roll (Roll) the getShape function from ‘DNAshapeR’ package was used ^44^. Input FASTA files, containing sequences in close proximity to identified DSB (±5nt or ±100nt), were extracted with custom python script (available upon request). The interaction potential (electrostatic and van der Waals) of Na^+^ probes with DNA duplexes was determined using a linear approximation to the Poisson-Boltzmann equation and dielectric constant for the DNA as implemented in the CMIP program ^45^.

### K-means clustering of DNA shape profiles

We used an elbow method to find appropriate number of clusters in the dataset, which consisted of predicted values of all parameters (MGW, HelT, ProT, Roll, EP) ±8 nt from each breakpoint. Based on cluster number diagnostic it was chosen to use k=9. Initial cluster centers were defined using 100 iterations. Next, we assigned every set of observations for each breakpoint into the closest centroid of 1 out of 9 clusters, independently for both – pks+ and pks-E.coli-induced DSB. Finally, sequence content of each cluster was exported and used as an input for computing proportion of each nucleotide per position (see SeqLogo method).

### SeqLogo

To compute and visualize the proportion of each nucleotide per position from collection of sequences consensusMatrix() and seqLogo() functions from ‘seqLogo’ package were used ^46,47^

### Model and Molecular dynamics set up

The 3D structure and protonation state of the colibactin were built starting from the smile (https://pubchem.ncbi.nlm.nih.gov/compound/138805674#section=InChI) using MarvinSketch (MarvinSketch, version 6.2.2, calculation module developed by ChemAxon, http://www.chemaxon.com/products/marvin/marvinsketch/). The geometry of the model and the partial atomic charges were assigned to the structure with General Amber Force Field (GAFF) ^48^. Parameters and topology files were prepared with Acpype ^49^. The colibactin was then simulated in explicit solvent at 298K (see below for details) for 250ns and along the simulation the distance between the cyclopropanes was monitored (see Fig.S4), to study their orientation and the overall length of the free colibactin. Using HADDOCK 2.4 ^16^, we then built the complex DNA-colibactin. For the docking, we selected a representative structure of the free colibactin along the MD simulation, with an average distance among the cyclopropanes (red line, Fig. S4) and an equilibrated structure of the DNA (sequence CGAAATTTCG). After the initial docking, that positioned the molecule correctly along the minor groove of the DNA, we then manually rotated slightly the molecule to improve the orientation of the cyclopropanes towards the N3 of the closest adenine using PYMOL (The PyMOL Molecular Graphics System, Schrödinger, LLC (2018)). To check the stability of this complex and to equilibrate its structure the model was simulated (see details MD simulation below) and minimized in solution with positional restraints on the solute using our well-established multi-step protocol ^50,51^. The minimized structure was thermalized to 298K at NVT, and then simulated first applying harmonic restraints of 5 kcal/mol·Å2 on the DNA on the DNA structure and distance constraints between the cyclopropane and the N3 of the adenine (respectively 4 and 5 bases apart), each represented by a harmonic restraint of 2.5 kcal/mol·Å^2^. To further check the stability of the complex we then slowly removed the constraints and run MD simulation of the complex during 60 ns by means of Molecular Dynamics simulations at NPT (P = 1 atm; T= 298K). The first 10 ns of the simulations were considered as an equilibration step and were discarded for further analysis.

In each MD simulation, DNA, free colibactin and their complex, respectively, we placed the solute in the centre of a truncated octahedral box of TIP3P water molecules ^52^, neutralized by K+ ions. In each simulation the Berendsen algorithm ^53^ was used to control the temperature and the pressure, with a coupling constant of 5 ps; and the SHAKE algorithm was utilized to equilibrium the length of hydrogen atoms involved in the covalent bonds ^54^. Long-range electrostatic interactions were accounted for by using the Particle Mesh Ewald method (14) with standard defaults, and a real-space cut-off of 10 Å. For the DNA we used the newly revised force field parmBSC1 ^55^. All simulations were carried out using AMBER 18 ^56^, and analyzed with CPPTRAJ ^57^ and visualized using VMD 1.9.4 ^58^.

### Cancer somatic mutation data

We obtained somatic variant data from the TCGA Unified Ensemble “MC3” Call Set ^59^ (“TCGA pan-cancer dataset”) and from the supplementary data of Giannakis et al ^18^. To test for enrichment of mutations at any motif we first identified positions of all hexanucleotide motifs in the exonic portion of the genome. Somatic variants occurring at A or T bases were grouped in one of 6 classes (quartile 1-4, outlier or POLE mutated sample) depending on the total SNV number and POLE mutation status of the corresponding tumor sample We then computed the mutation rate for each hexanucleotide motif with respect to the number of genomic bases covered in exonic regions for the same motif. As a baseline, we established the mutation rates of all WWWWWW motifs and subtracted their mean from the mutation rate of all other hexanucleotide motifs. We then tested for significance of the mutation rate at colibactin associated AAWWTT motifs (i.e. AAATTT and AAAATT/AATTTT) compared to the remaining WWWWWW motifs using Mann-Whitney-U tests and computed the false discovery rate (FDR) using the method of Benjamini-Hochberg ^60^. Reads from WGS of colorectal cancers ^19^ EGA database accession code <u>EGAS00001003010</u>,) were aligned to GRCh38 with BWA-MEM ^41^ and called using Mutect2 ^61^. All single nucleotide variant calls (PASSed by Mutect2) were used to determine the number of mutations overlapping WWWWW pentanucleotides and WWWWWW hexanucleotides and further analyzed in a similar way as for exome sequencing data on an individual sample basis.

### Analysis of pattern enrichment in cancers

For analysis of signatures we classified all variants according to the presence of patterns in the +/-5bp around SNV variant calls: one group contained colibactin associated pentanucleotides (AAATT/AATTT or AAAAT/ATTTT), one contained AAA/TTT in order to control for AT-rich sequences and one contained all other motifs. The R package deconstructSigs ^62^ was used to estimate the contribution of COSMIC signatures v3 ^21^ independently for each group. Differences between groups were assessed for each single base change signature (SBS) between groups using Mann-Whitney test.

### Data analysis and visualization

All visualizations and statistical analyses were produced using R v3.4 ^63^

## Acknowledgements

The results shown here are in whole or part based upon data generated by the TCGA Research Network: https://www.cancer.gov/tcga. The authors would like to thank Dr. Silvano Garnerone from N.C. laboratory for processing raw BLISS data, Prof. Pablo D. Dans from Biophysical Chemistry Lab., Department of Biological Science (CENUR Litoral Norte), UdelaR, UY, for constructive discussions about the theoretical model of colibactin, Prof. Ulrich Dobrindt from University of Münster for providing E.coli strains and Rike Zietlow for editing the manuscript.

## Funding

P.J.D.K. was supported by IMPRS. B.A.M.B was supported by a Rubicon fellowship from the Netherlands Organisation for Scientific Research (NWO). M.O. is an ICREA (Institució Catalana de Recerca i Estudis Avancats) academia researcher. This work was supported by the Spanish Ministry of Science (grants BIO2015-64802-R-), Spanish Ministry of Science (BFU2014-61670-EXP and BFU2014-52864-R), the Catalan Government (grants 2014-SGR); the Instituto de Salud Carlos III-Instituto Nacional de Bioinformática (ISCIII PT 13/000/0030 co-funded by the Fondo Europeo de Desarrollo Regional [FEDER] and the Biomolecular and Bioinformatics Resources Platform) and the European Union’s Horizon 2020 research and innovation program (grants Elixir-Excelerate: 676559; BioExcel2:823830), the MINECO Severo Ochoa Award of Excellence from the Government of Spain (awarded to IRB Barcelona), the Karolinska Institutet, the Ragnar Söderberg Foundation, the Swedish Foundation for Strategic Research (to N.C.: BD15-0095), and the Strategic Research Programme in Cancer (StratCan) at Karolinska Institutet.

## Author contributions

P.J.D.K, H.B. and T.F.M. designed experiments, P.J.D.K. and B.A.M.B. performed sBLISS experiments, A.I. performed E.coli infection and immunostaining. Bioinformatics analysis were performed by P.J.D.K. and H.B. Theoretical model of colibactin was built by F.B. and M.O. R.K. and L.A.A. provided and analyzed WGS colorectal cancer data. The manuscript was written by P.J.D.K., H.B., F.B. and T.F.M.

## Competing interests

The authors declare no competing interests

## Data and materials availability

Input FASTA files and analysis scripts are available upon request. All other data is available in the main text or supplementary materials.

## Supplementary Materials

### Supplementary Figure Legends

**Fig. S1.**
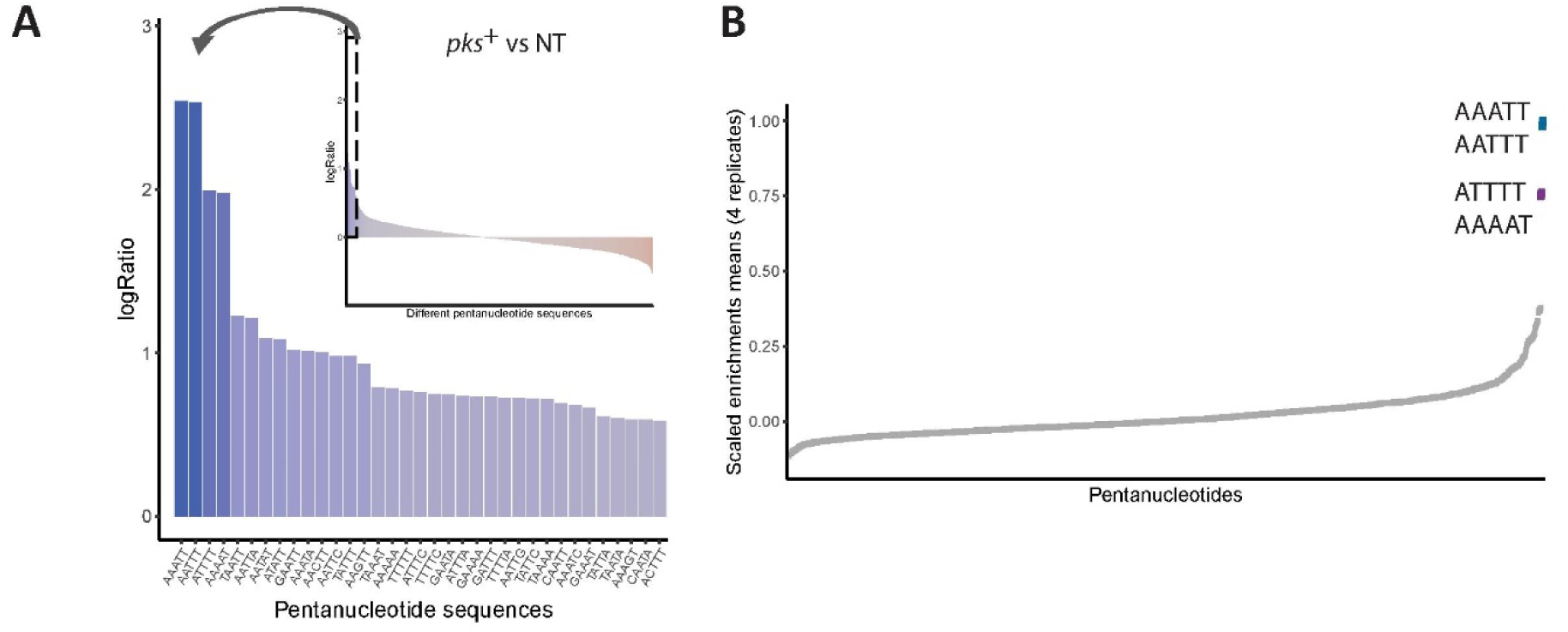
Outstanding enrichment of AAATT and ATTTT and their reverse and complement sequences in colibactin-induced DSBs. (A) Pentanucleotide sequences enriched (log2 ratio of proportions of DSB at each motif between both conditions) at the DSB positions caused by colibactin-positive E.coli (pks+) in comparison to non-treated (NT) cells. (B) Scaled enrichment means of all 4 independent biological replicates for all possible pentanucleotides (1,024) obtained by comparing nucleotide content of pks+ E.coli infection-induced breaks to the pks-E.coli-induced breaks.

**Fig. S2.**
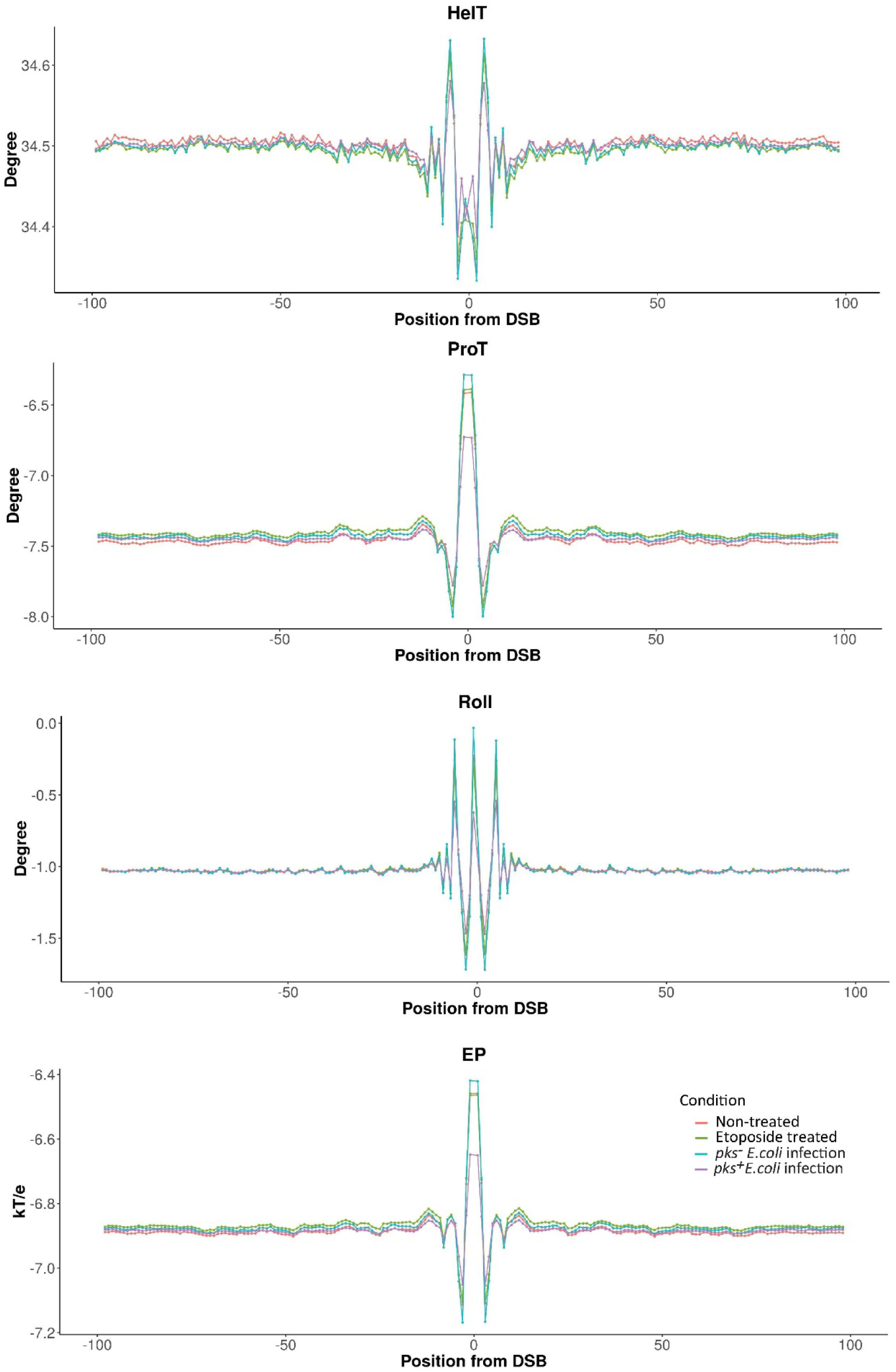
Averaged values of DNA shape properties (HelT, ProT, Roll and EP) for sequences in proximity to identified DSBs of non-treated, etoposide-treated, pks-E.coli infected and pks+ E.coli infected cells.

**Fig. S3.**
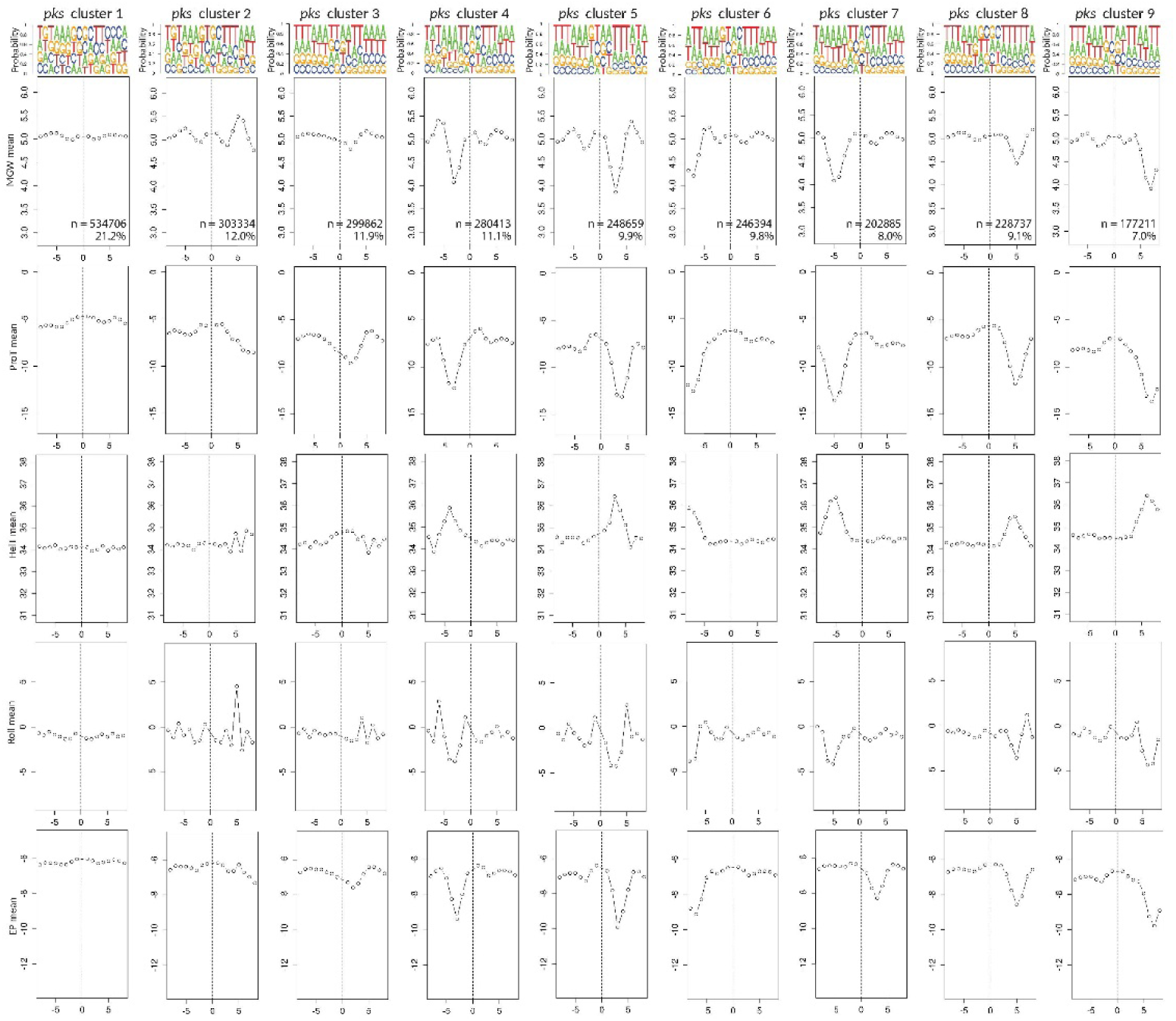
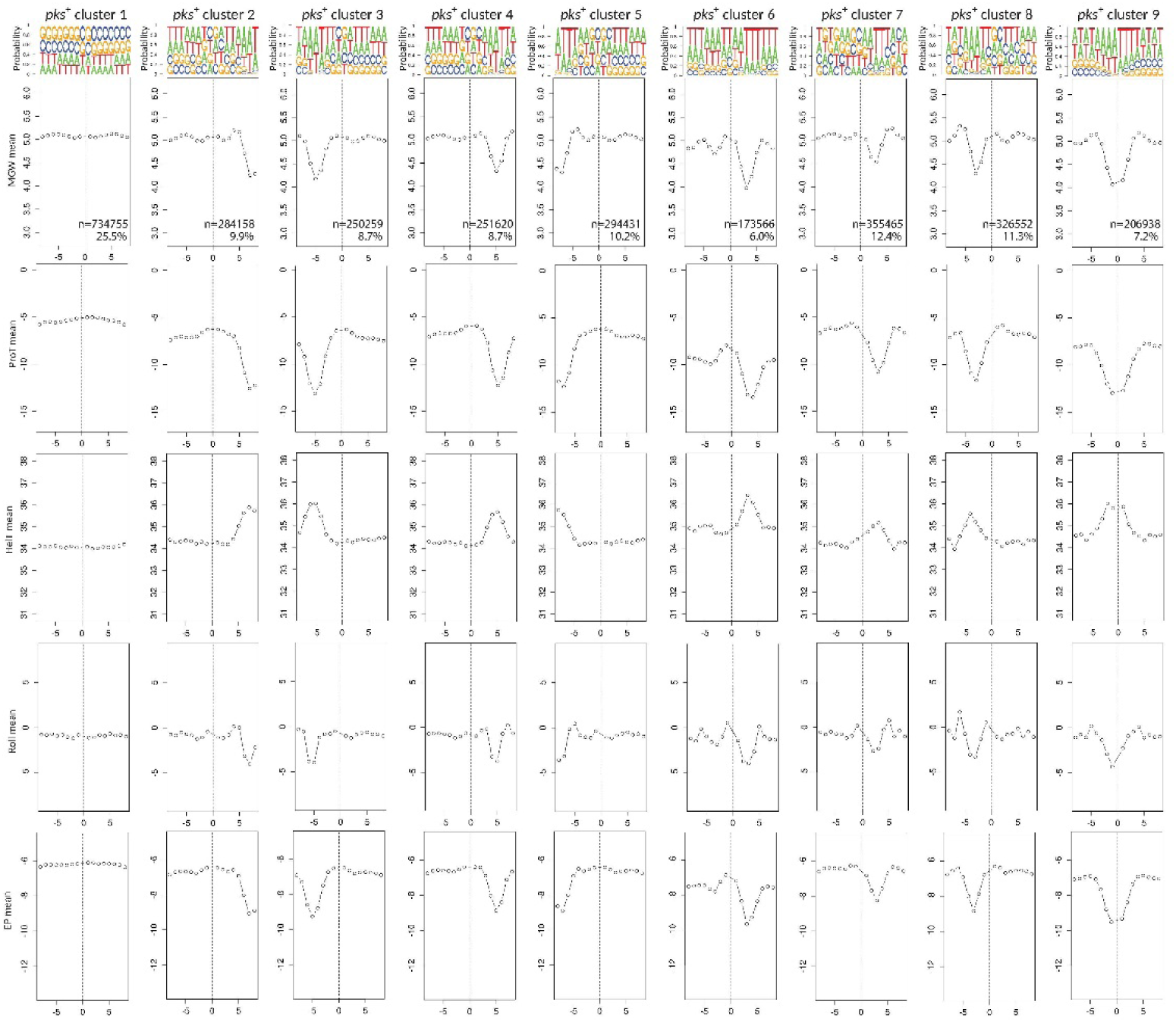
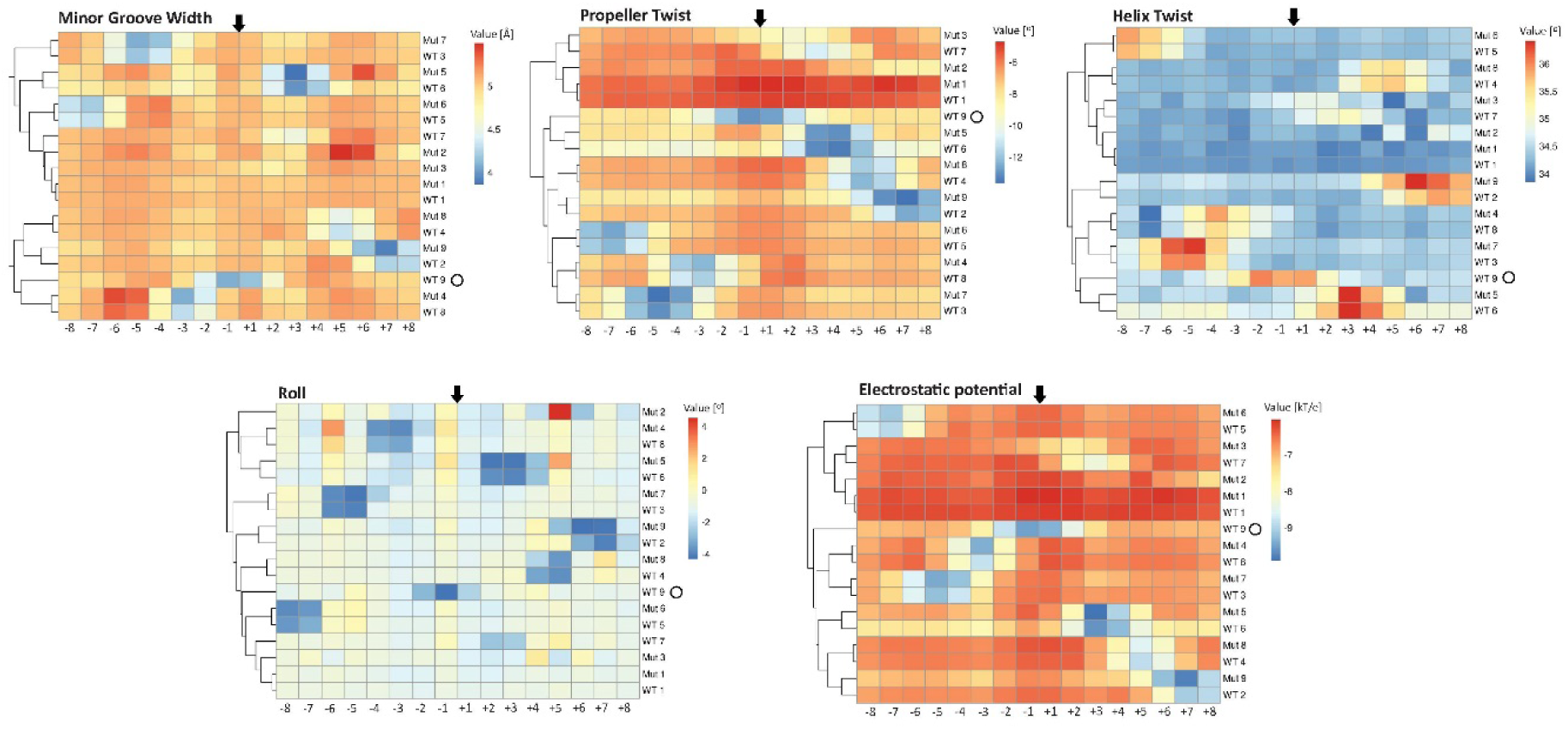
K-means clustering of all predicted values of the DNA shape parameters. (A-B) DNA shape profiles of all clusters obtained from k-means clustering of pks+ and pks-conditions. (A) Clusters identified form the pks-dataset. (B) Clusters identified from the pks+ dataset. Above each cluster nucleotide probability for every position is presented. (C) Heatmaps comparing averaged profiles of all identified clusters in pks+ and pks-conditions based on predicted DNA shape parameters, presented as individual comparisons of averaged profiles for each DNA shape parameter. Colors indicate absolute values for each DNA shape characteristics. Each square in the heatmap refers to specific position from the break. Black arrows are marking exact DNA DSB position.

**Fig. S4.**
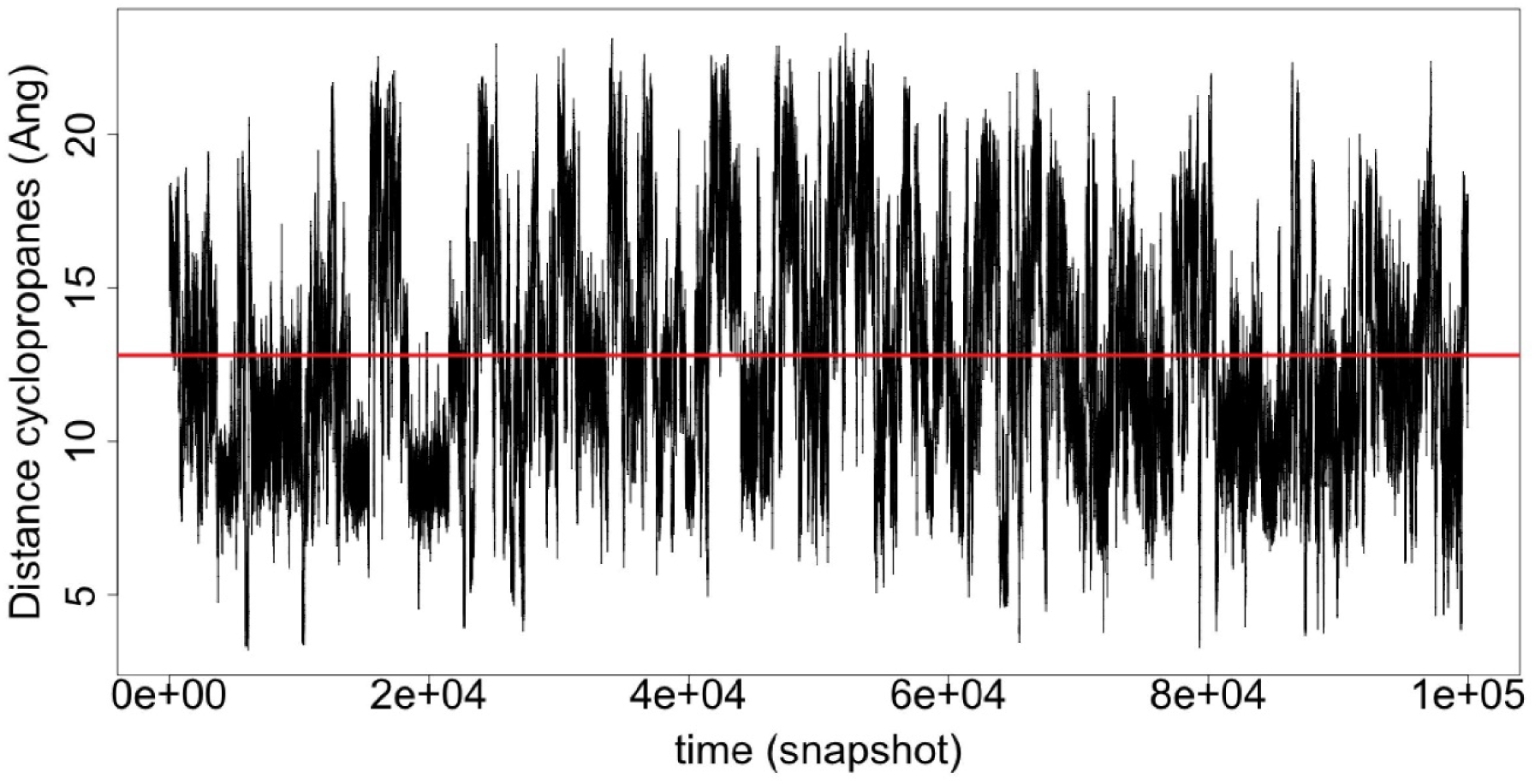
Distance of the cyclopropanes (Å) along the MD simulation of the free colibactin in water. Red line identifies the average values of this distance (12.8Å).

**Table S1** Summary of DSB enrichment at all 1024 pentanucleotide patterns across 4 replicates with associated predicted central DNA shape characteristics corresponding to Fig. 2A/B and Fig. 4A

